# Anaerobic methane oxidation by ANME-2a at 2 M chloride in the Orca Basin halocline

**DOI:** 10.64898/2026.02.09.704686

**Authors:** Lea A. Adepoju, Emily R. Paris, Jordan M. McKaig, Rebecca S. R. Salcedo, Aaron Martinez, Steffen Buessecker, Emilie J. Skoog, Claire E. Elbon, Miguel Desmarais, Cathryn D. Sephus, Chad I. Pozarycki, Veronica Hegelein, Enrica S. Quartini, Amina T. Schartup, Peter T. Doran, Christopher E. Carr, Ellery D. Ingall, Douglas H. Bartlett, Ranjani Murali, Morgan R. Raven, Britney E. Schmidt, Jeff S. Bowman, Anne E. Dekas, Jennifer B. Glass

**Author notes:** Corresponding author: Jennifer B. Glass.

## Abstract

Anaerobic methane oxidation, typically mediated by consortia of archaea and bacteria, is a key process in the global methane cycle, but little is known about its upper salinity limits. We characterized the microbial methane cycle in the anoxic, hypersaline Orca Basin using metagenomics, metatranscriptomics, fluorescence *in situ* hybridization, and geochemical measurements at sub-meter resolution. In the brine, we detected transcriptional activity of the halophilic methylotrophic methanogen *Methanohalophilus*, consistent with a biological source for Orca Basin methane. In the particle-rich halocline (∼2 M Cl^-^; ∼2235 meters depth), high *mcrA* transcription by a novel ANME-2a species was co-located with a positive shift in δ^13^C-CH_4_ consistent with anaerobic oxidation of methane. ANME-2a also transcribed genes for biosynthesis of the osmolyte N(ε)-acetyl-β-L-lysine, supporting adaptation to hypersaline conditions. At the same depth, consortia of sarcina-like archaea, likely ANME-2a, were observed in association with vibrioid and filamentous bacteria. Members of a halotolerant Desulfobulbales genus (family SURF-16, which includes the previously identified ANME partner Seep-DBB) were active at the same depth. At and above the oxic-anoxic interface, aerobic methane oxidation appears to be mediated by three genera of uncultivated Methylococcales bacteria. Our results extend the known upper salinity range of ANME-2a to ∼2 M Cl^-^ and reveal the key microbial players in the methane bio-filter between the Orca Basin brine and overlying seawater.

## Introduction

Orca Basin is a deep hypersaline anoxic basin (DHAB) in the Gulf of Mexico formed by dissolution of Jurassic-age halite deposits over the past 8,000 years [1, 2]. High concentrations of biogenic methane (∼700-900 μM) are present in Orca Basin’s NaCl-saturated brine and sediments [3, 4]. Methanogenesis in Orca Basin [5, 6] and Mediterranean and Red Sea DHABs [7–15] is likely mediated by halophilic methylotrophic *Methanohalophilus*. Despite abundant sulfate, anaerobic methane oxidation (AOM) has not previously been detected in Orca Basin brine and sediments [5, 6, 16], potentially due to energetic limitation under salt stress [17]. There is geochemical [18, 19] and microbial [15, 20–22] evidence for methane oxidation in the sharp haloclines (1-3 meters) at the brine-seawater interfaces in Red Sea and Eastern Mediterranean DHABs, but little is known about methanotrophy in the expansive (∼80 meter) Orca Basin chemocline. In this study, sampling at sub-meter depth resolution across the Orca Basin chemocline allowed us to pinpoint the precise location of anaerobic and aerobic methane oxidation and to identify the microbes mediating these processes by combining metatranscriptomics, microscopy, and isotope geochemistry.

## Results and Discussion

### Geochemical gradients and methane dynamics in Orca Basin

Eight sites (P1, P2, P5 in the North Orca Basin and P8, P9, P13, P14, and P15 in South Orca Basin) were sampled in 2023 as described by Paris et al [23] **(Fig. S1**). Geochemical profiles for the North Orca Basin sites (P1, P2 and P5) were previously reported by Paris et al [23]. Depth profiles were remarkably consistent across the basin. In the halocline (2235-2250 meters water depth), chloride (Cl^-^) concentration rose from 1 to 4-5 M **(Table S1; Fig. 1a)**, salinity, measured as total dissolved solids, increased from ∼80 to ∼240 ppt **(Table S1)**, and water activity dropped from 0.96 to 0.78 a_w_ (**Table S1)**. Temperatures increased from 4.3°C at 2200 meters to 5.6°C in the brine **(Table S1)**. pH was highest (∼8.2) in the halocline and lowest (∼6.5) in the brine **(Table S1).** A prominent nepheloid particle layer [24] in the halocline was apparent in the drop in beam transmission **(Fig. 1b)**. The oxycline extended from 2170 m to 2200 meters, and dissolved O_2_ was undetectable below 2201 meters **(Table S1; Fig. 1c)**. Nitrate concentrations dropped through the oxycline and were undetectable below 2212 meters **(Table S1; Fig. 1d)**. Dissolved Mn^2+^ began to increase at 2201 meters, displayed a prominent peak (410 μM) in the halocline at ∼2235 meters, and was present at ∼150 μM in the brine **(Table S1; Fig. 1e)**, consistent with a previous study [25]. Dissolved Fe^2+^ rose at the halocline-brine interface and was present at ∼40 μM below 2250 meters (**Table S1; Fig. 1f)**, also consistent with a previous study [25]. Sulfate rose from seawater concentrations (∼28 mM) to ∼50 mM across the halocline **(Table S1; Fig. 1g).** Sulfide was below the detection limit of ∼1 μM throughout the water column. Methane was ∼700-900 μM in the brine, ∼5-20 μM in the chemocline, and ∼0.001-0.005 μM in the overlying seawater (**Table S1**; **Fig. 1h**; Wiesenburg et al [3]). Brine samples in 2023 had δ^13^C-CH_4_ values of -76 ‰ **(Table S1; Fig. 1i)**, consistent with previous studies [3, 4]. In the halocline, δ¹³C-CH increased to -68 ‰ (**Table S1; Fig. 1i)**, likely as a result of isotope fractionation during AOM [26], although it is possible that the isotopic shift could result from mixing or other processes.

**Figure 1.**
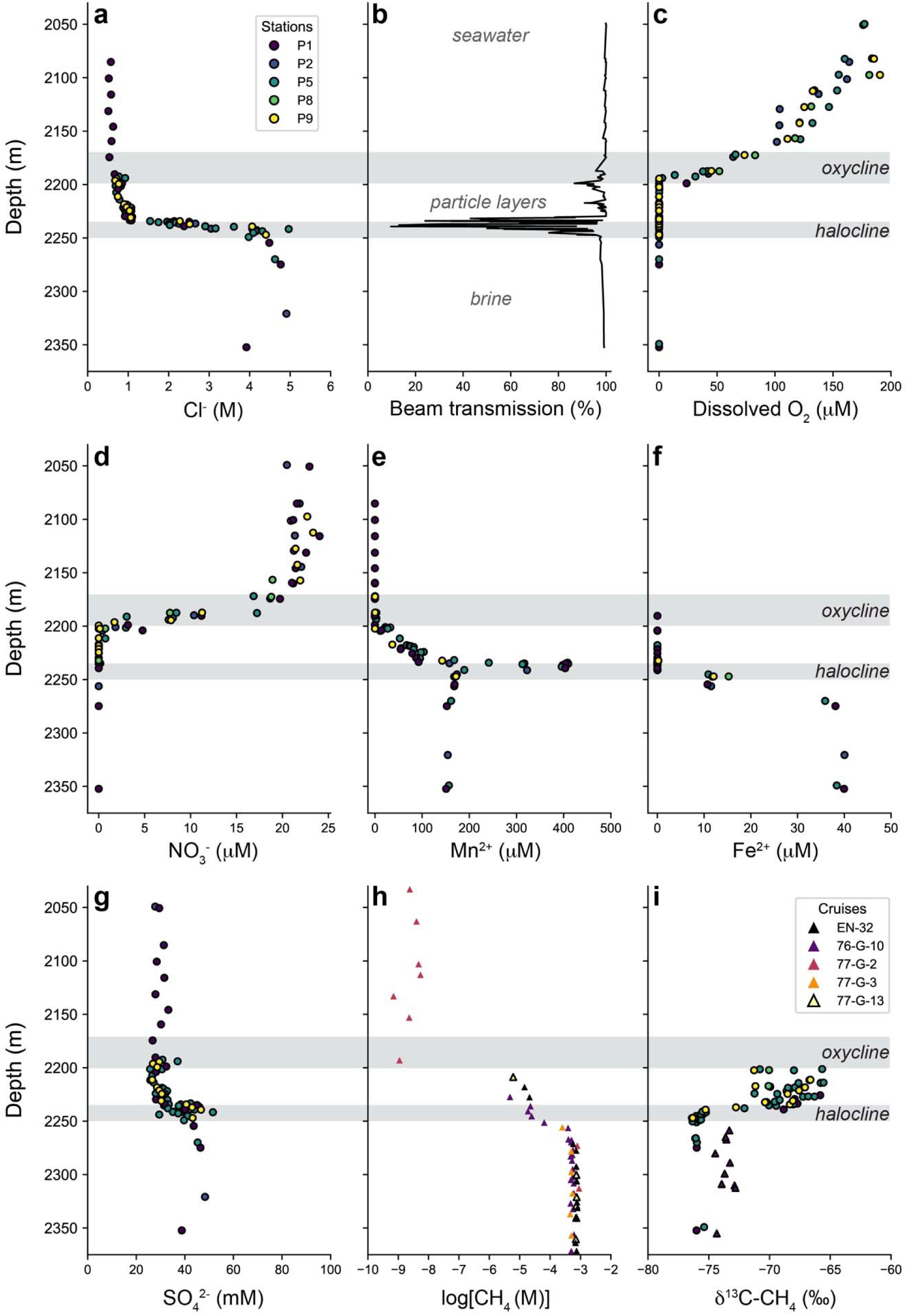
Depth profiles of key geochemical parameters through the Orca Basin water column. (a) Chloride (Cl^-^) concentration; (b) beam transmission (%) (from stations P1-P9); (c) dissolved oxygen (O_2_) concentration; (d) nitrate (NO ^-^) concentration; (e) dissolved manganese (Mn^2+^) concentration; (f) dissolved iron (Fe^2+^) concentration; (g) dissolved sulfate (SO ^2-^) concentration; (h) methane concentration from Wiesenburg et al [3]; (i) δ^13^C-CH_4_ isotopic composition from Wiesenburg et al [3] (triangles) and the present study (circles). The upper gray box indicates the depth range of oxycline (2170-2200 meters) and the lower gray box indicates the depth range of the halocline (2235-2250 meters). Data in panels a, b, c, d, e, f, and g for stations P1, P2, and P5 were previously reported by Paris et al [23].

The close agreement between our 2023 measurements and those of Weisenburg, Brooks, and Bernard (collected ∼40 years earlier) in methane concentration, δ^13^C-CH_4_, and salinity profiles corroborates previous findings [27] that the geochemistry of the Orca Basin brine is stable over decadal timescales. This stability reflects the physically isolated nature of the basin and its slow hydrological exchange with overlying seawater and provides the sustained environmental conditions necessary for the establishment of slow-growing microbial communities.

### *Methanohalophilus* is transcriptionally active in Orca Basin brine

The methylotrophic methanogenic archaeon *Methanohalophilus* was present at low abundance (0.01-0.05% total amplicon sequence variants, or ASVs) in the Orca Basin brine **(Table S1)**, consistent with the similarly low relative abundance of *Methanohalophilus* in Orca Basin sediments [6]. One metagenome assembled genome (MAG 231106-OB-CTD-1-B3-DNA_23; 56% estimated completeness; **Table S2**) lacking 16S rRNA and *mcrA* genes was assigned to an unknown species in the genus *Methanohalophilus* (91.3% ANI to *Methanohalophilus halophilus* Z-7982; **Table S3**). Transcription of tetrahydromethanopterin S-methyltransferase subunit A (*mtrA*) increased with depth in the brine to ∼20 transcripts per million (TPM) at 2309 m **(Table S1)**. *Methanohalophilus* transcription in the brine, coinciding with highly negative δ^13^C-CH_4_ values, supports previous findings that the methane in the Orca Basin brine is from microbial sources [3, 6] and that biological methylotrophic methanogenesis is active in Orca Basin, as in other DHABs [7–12]. The sharp increase in *Methanohalophilus* abundance and *mtrA* transcription at the halocline-brine interface also coincides with the steep increase in salinity, consistent with the required higher salinity optimum conditions of this genus [14] and suggest that methanogenesis is largely restricted to the NaCl-saturated brine.

### ANME-2a is transcriptionally active in Orca Basin halocline

Metagenomic and transcriptomic data from the deep brine do not reveal clear evidence of AOM activity. In contrast, the increase in δ¹³C-CH values (**Table S1; Fig. 1i)** in the halocline, where dissolved O_2_ is under detection, is suggestive of AOM. We recovered one ANME-2a (*Methanocomedens* [28]) MAG (231106-OB-CTD-1-B3-DNA_filter_8; 93% estimated completeness; **Table S2**) and one ANME-2d MAG (*Methanoperedens_*A; 231107-OB-CTD-2-B5-DNA_65; 73% estimated completeness; **Table S2**). The ANME-2a MAG contained a 16S rRNA gene that 100% matched the most abundant ANME ASV, which peaked at and just under the halocline (∼0.1% total ASVs; **Tables S1, S2; Fig. 2a)**. Transcription of ANME-2a *mcrA* was extremely high in the halocline (∼2500 TPM; **Table S1; Fig. 2b)**, at the same depth as the positive shift in δ^13^C-CH_4_ **(Fig. 1f)**, relative to the rest of the water column. The phylogeny of concatenated single-copy archaeal proteins showed that the Orca Basin ANME-2a belongs to a different GTDB species than existing MAGs from marine sediment cold seeps **(Fig. 2c)**.

**Figure 2.**
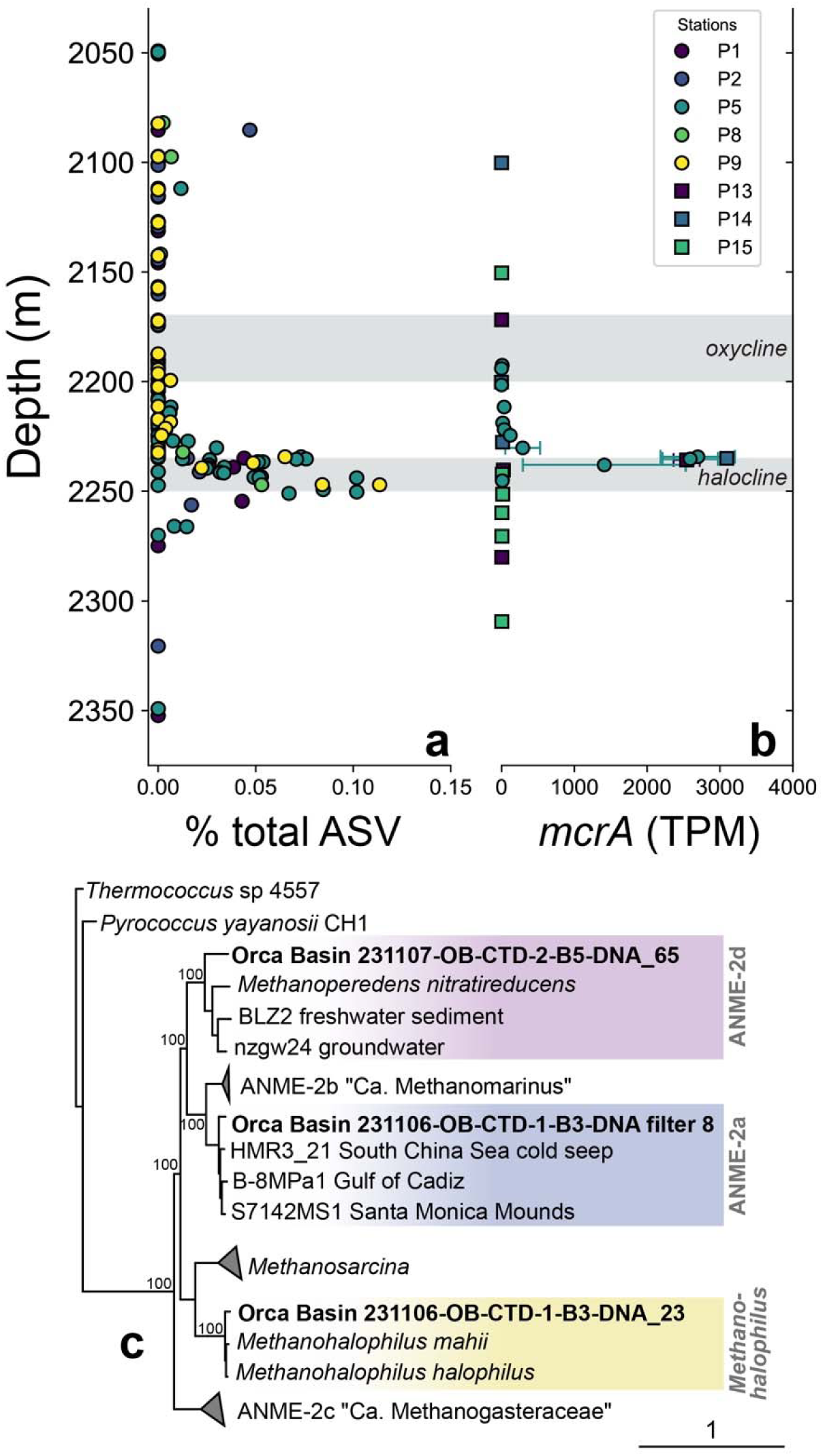
Relative abundance and *mcrA* transcription of ANME-2a through the Orca Basin water column, and Methanosarcinales phylogeny showing genera present in Orca Basin (colored bars). (a) Relative abundance (ASV %); (b) *mcrA* gene transcription (TPM); (c) concatenated single copy marker protein phylogeny of select members of the Methanosarcinales order (ANME-2a-d, *Methanosarcina,* and *Methanohalophilus*), with Orca Basin MAGs in bold. In panels a and b, the upper gray box indicates the depth range of oxycline (2170-2200 meters) and the lower gray box indicates the depth range of the halocline (2235-2250 meters).

The high activity of ANME-2a in the Orca Basin halocline (2 M Cl^-^) is notable because ANME-2a are rarely reported in hypersaline environments [29, 30]. To our knowledge, the highest Cl^-^ concentration with prevalence of ANME-2a is 0.8 M in Napoli mud volcano hypersaline sediments [31]; at 1.2 M Cl^-^, ANME-1 was more prevalent than ANME-2a [31]. The Orca Basin halocline may represent a rare exception where ANME-2a activity is confined to a narrow depth interval in the halocline and is absent from the underlying methane and sulfate-rich brine, potentially implicating salinity as an important environmental control on AOM in the Orca Basin.

The likely mechanism for halotolerance in Orca Basin ANME-2a is biosynthesis of N(ε)-acetyl-β-L-lysine, the dominant osmolyte at 2 M osmolality in *Methanosarcina* spp. [32], which is also synthesized by ANME-2d under salt stress [33]. N(ε)-acetyl-β-L-lysine is synthesized by lysine-2,3-aminomutase (encoded by *kamA*) and β-lysine-N6-acetyl transferase (encoded by *ablB*), both of which were transcribed by ANME-2a in the halocline (**Table S1).** The *kamA-ablB* genes are adjacent on a ten-gene contig; the eight other genes had 81-94% identity to ANME-2a MAGs, whereas the *kamA* and *ablB* genes had highest identity (62% and 82%, respectively) to ANME-2d MAGs **(Table S4)** and were likely acquired by horizontal gene transfer [33]. The Orca Basin ANME-2a also transcribed a multiheme (37) cytochrome (MHC) S-layer domain protein (**Table S1, S5)**, similar to those previously described in other ANME-2 for extracellular electron transfer to either a bacterium or an electron shuttle [34, 35].

Other ANME lineages (ANME-2d (*Ca.* Methanoperedens) and ANME-1b) were present throughout the chemocline at very low abundance (0.001-0.005% total ASVs; **Table S1)**. The ANME-2d MAG contained six weakly transcribed MHCs (12-25 hemes; ∼4 TPM at ∼2235 meters **(Tables S1, S4)**. The presence of ANME-2d in the water column is notable, as this lineage is rarely reported in deep-sea settings [36, 37], and is known to couple AOM with reduction of nitrate [38], manganese [39], and iron oxides [40]. The prominent Mn^2+^ peak in the Orca Basin halocline **(Fig. 1e)** raises the possibility that manganese could also serve as an electron acceptor for AOM in Orca Basin, either directly or indirectly [41]. Furthermore, the presence of Fe(OH)_3_ in the halocline has been proposed as an abiotic product of Fe^2+^ and manganese oxides [25], raising the possibility that Fe-AOM may also be a contributing pathway.

### Archaeal-bacterial consortia in Orca Basin halocline revealed by CARD-FISH

Catalyzed reporter deposition-fluorescence *in situ* hybridization (CARD-FISH) using ARCH915 (archaea) and EUB338 (bacteria) revealed consortia of sarcina-like archaea, often partially associated with vibrioid and filamentous bacteria, in halocline samples (3.4 x 10^4^ consortia mL^-1^; **Fig. 3a-d; Figs. S2-S6; Table S6)** from 2234 meters (2.1 M Cl^-^). The observed archaeal-bacterial consortia were 2-6 μm diameter and contained 1-4 bacterial cells and 1-9 archaeal cells each (n=14). We hypothesize that the archaea with sarcina-like morphology are ANME-2a, as the only other archaeal 16S rRNA ASVs at 2234 meters belong to Nitrososphaeria, which are not known to form sarcina morphologies or physical associations with bacteria. Confirmation of archaeal identity awaits future work with validated probes. In some of the consortia, bacterial cells appeared to partially envelope or curve around the archaeal clusters (**Fig. 3b,c**), while in others, bacteria appeared to extend outward from archaeal cluster surfaces (**Fig. 3d**). The morphology of the Orca Basin archaea-bacteria consortia is often similar to the “partial shell” association commonly observed for ANME-bacteria consortia from methane seep sediments, including for ANME-Desulfobulbales (DBB) [42, 43].

**Figure 3.**
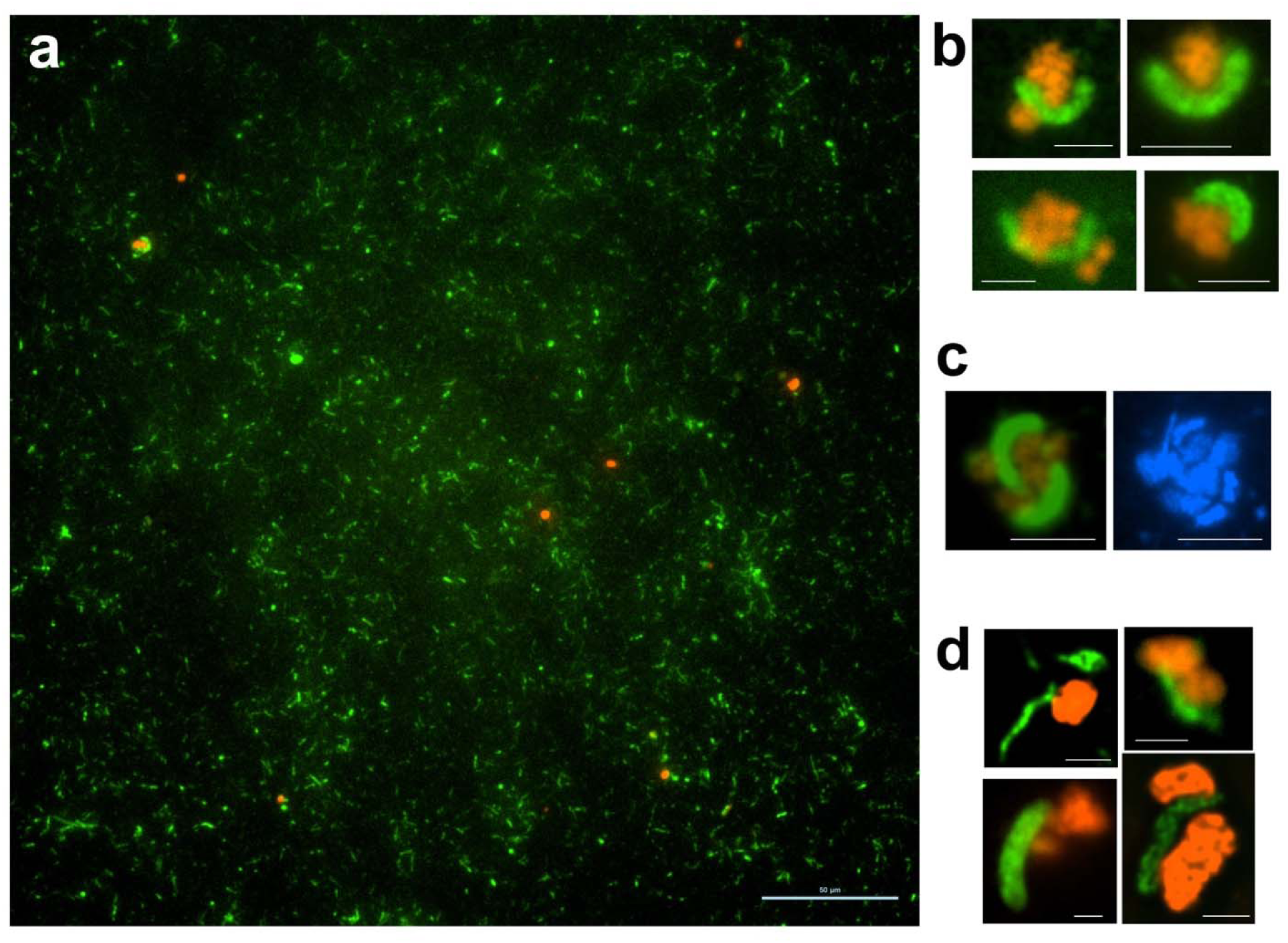
CARD-FISH microscopy of archaea-bacterial consortia in samples from 2234 meters depth in the Orca Basin halocline with probes ARCH915 (archaea; red) and EUB338 (bacteria; green). Scale bar: 50 μm for panel a, 5 μm for panels b and c, and 2 μm for panel d. Panel c shows the FISH image on the left and the corresponding DAPI image (blue) on the right.

### Halophilic SURF-16/Seep-DBB sulfate-reducing bacteria peak in activity and abundance in the halocline

ANME-2a is best known to associate with Seep-SRB1a bacteria (GTDB taxonomy: p Desulfobacterota; c Desulfobacteria;o Desulfobacterales; f ETH-SRB1; g B13-G4 [44, 45]) but can be associated with other bacterial partners [46, 47]. No high-quality ETH-SRB1 MAGs were present in our dataset, and only one ASV classified as Seep-SRB1 was present at low abundance (0.01%) in some, but not all, halocline samples **(Table S1).** Instead, we found that sulfate-reducing bacteria with GTDB taxonomy p Desulfobacterota; c Desulfobulbia; o Desulfobulbales; f SURF-16; g JAWLDV01 peaked in relative abundance **(Fig. 4a)** and *dsrA* transcription **(Fig. 4b)** at the same depth as ANME-2a (2234-2235 meters; **Table S1)**. JAWLDV01 was approximately eight times more abundant than ANME-2a at 2234-2235 meters based on ASV relative abundance **(Table S1)**, suggesting that the majority of the JAWLDV01 population is likely free-living [5, 48]. However, variable cell lysis efficiency of bacterial and archaeal cells during nucleic acid extraction could also affect the observed relative abundance of these groups.

**Figure 4.**
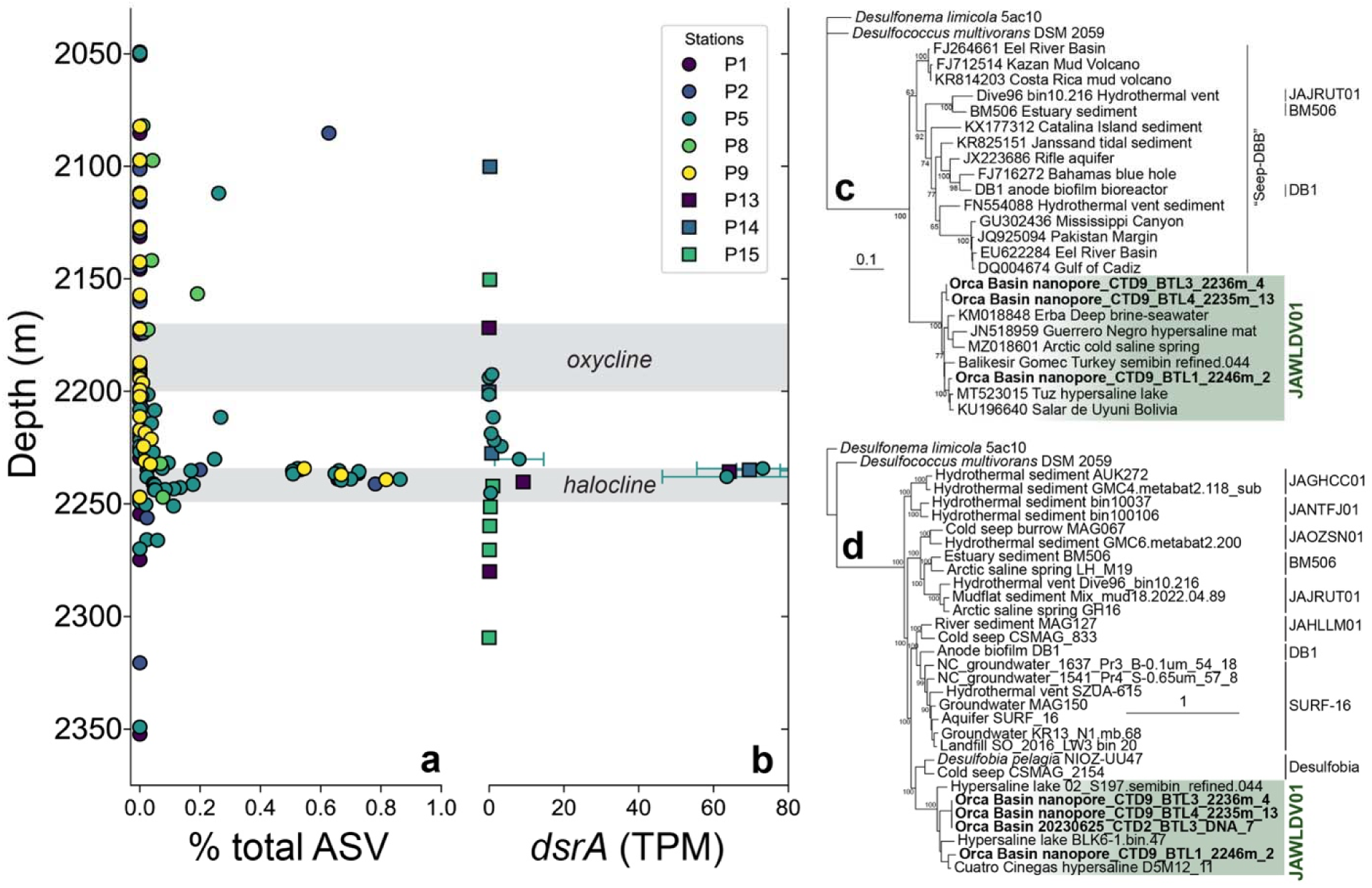
Relative abundance and *dsrA* transcription of genus JAWLDV01 (MAG nanopore_CTD9_BTL3_2236m_4) through the Orca Basin water column, and phylogenies of family SURF-16 (order Desulfobulbales). (a) Relative abundance (ASV %); (b) *dsrA* gene transcription (TPM); (c) 16S rRNA phylogeny for family SURF-16, including those formerly known as “Seep-DBB”; (d) concatenated bacterial marker protein phylogeny for family SURF-16. In phylogenies, bold indicates the MAG used for metatranscriptomic mapping and labels are GTDB-R232 genera. In panels a and b, the upper gray box indicates the depth range of oxycline (2170-2200 meters) and the lower gray box indicates the depth range of the halocline (2235-2250 meters).

To investigate the phylogeny of JAWLDV01 sulfate-reducing bacteria in Orca Basin, we retrieved full length 16S rRNA sequences from two other replicate bins (nanopore_CTD9_BTL3_2236m_4 and nanopore_CTD9_BTL4_2235m_13; **Table S2**) as well as an additional MAG from 2246 meters (nanopore_CTD9_BTL1_2246m_2; **Table S2**), because the MAG with highest completeness (91%) and lowest contamination (0.2%; 20230625_CTD2_BTL3_DNA_7) used for metatranscriptomic mapping lacked a 16S rRNA gene. The 16S rRNA genes from these Orca Basin MAGs form a genus-level clade with other sequences from hypersaline environments **(Fig. 4c)**, suggesting that JAWLDV01 is a halophilic genus. The Orca Basin JAWLDV01 16S rRNA genes share 88-91% sequence similarity with Seep-DBB bacteria from methane seep sediments and mud volcanoes [43, 48] **(Fig. 4c)**, strongly suggesting that the Seep-DBB clade in AOM literature is either synonymous with SURF-16 in GTDB taxonomy, or a genus-level clade within that family. The SURF-16 family also includes MAGs from hydrothermal sediments and groundwater (**Fig. 4c, d)**.

ANME-2 have previously been reported in association with Seep-DBB [42, 43, 48] and other Desulfobulbales [47]. Desulfobulbales are also known to contain MHCs that have been hypothesized to be involved in extracellular electron transfer from ANME [45]. At 2234-2235 meters, the Orca Basin JAWLDV01 MAG transcribed the porin:decaheme cytochrome OmcKL found in syntrophic partners HotSeep-1, Seep-SRB2, and Seep-SRB1g (83 ± 8 TPM; **Table S5**) and a novel porin:decaheme cytochrome complex (24 ± 5 TPM; **Table S5**), consistent with a capacity for syntrophic electron transfer [45]. However, JAWLDV01 is unlikely to be an obligate syntrophic partner because it appears to also use hydrogen as an electron donor; the Orca Basin JAWLDV01 MAG transcribed genes for a membrane-bound nickel-iron hydrogenase (**Table S1)**, which is lacking in obligate syntrophic SRB, but are present in facultative syntrophic Desulfofervidales [45].

Whether JAWLDV01 represents a primary syntrophic partner or one of multiple bacterial partners, including iron-reducing bacteria [49–51], or is completely free-living, remains uncertain. ANME-2a are known to possess versatile electron transfer pathways [52] and can decouple from sulfate-reducing bacteria when provided an external electron shuttle [53, 54]. It is possible that the stable particle layer in the Orca Basin halocline enables electron shuttling and/or that metal oxides in the particle layer stimulate AOM by facilitating cryptic sulfur cycling [55, 56].

### Aerobic methanotrophs are transcriptionally active at and above the Orca Basin oxic-anoxic interface

Methane that escapes AOM in the Orca Basin halocline appears to be oxidized aerobically higher in the chemocline by gammaproteobacterial methanotrophs of the order Methylococcales, which were transcriptionally active at and above Orca Basin’s oxic-anoxic interface. The genera UBA1147 (family UBA1147; MAG 20230701_CTD10_BTL11_DNA_filter_6; 84.6% estimated completeness; **Table S2**) and OPU3-GD-OMZ (family Methylomonadaceae; MAG 231108-OB-CTD-3-B9-DNA_filter_49; 88.3% estimated completeness; **Table S2**) were present at 1-4% and 0.5-1.5% total ASVs, respectively **(Fig. 5a,c)** and transcribed *pmoA* at 2000-4000 TPM and 100-600 TPM, respectively (**Fig. 5b,d**), peaking at ∼2150 meters (100-130 µM dissolved O_2_; **Table S1)**. These genera have previously been implicated in methane oxidation in coastal regions (e.g. oxygen minimum zones and overlying methane cold seeps [57–59]).

**Figure 5.**
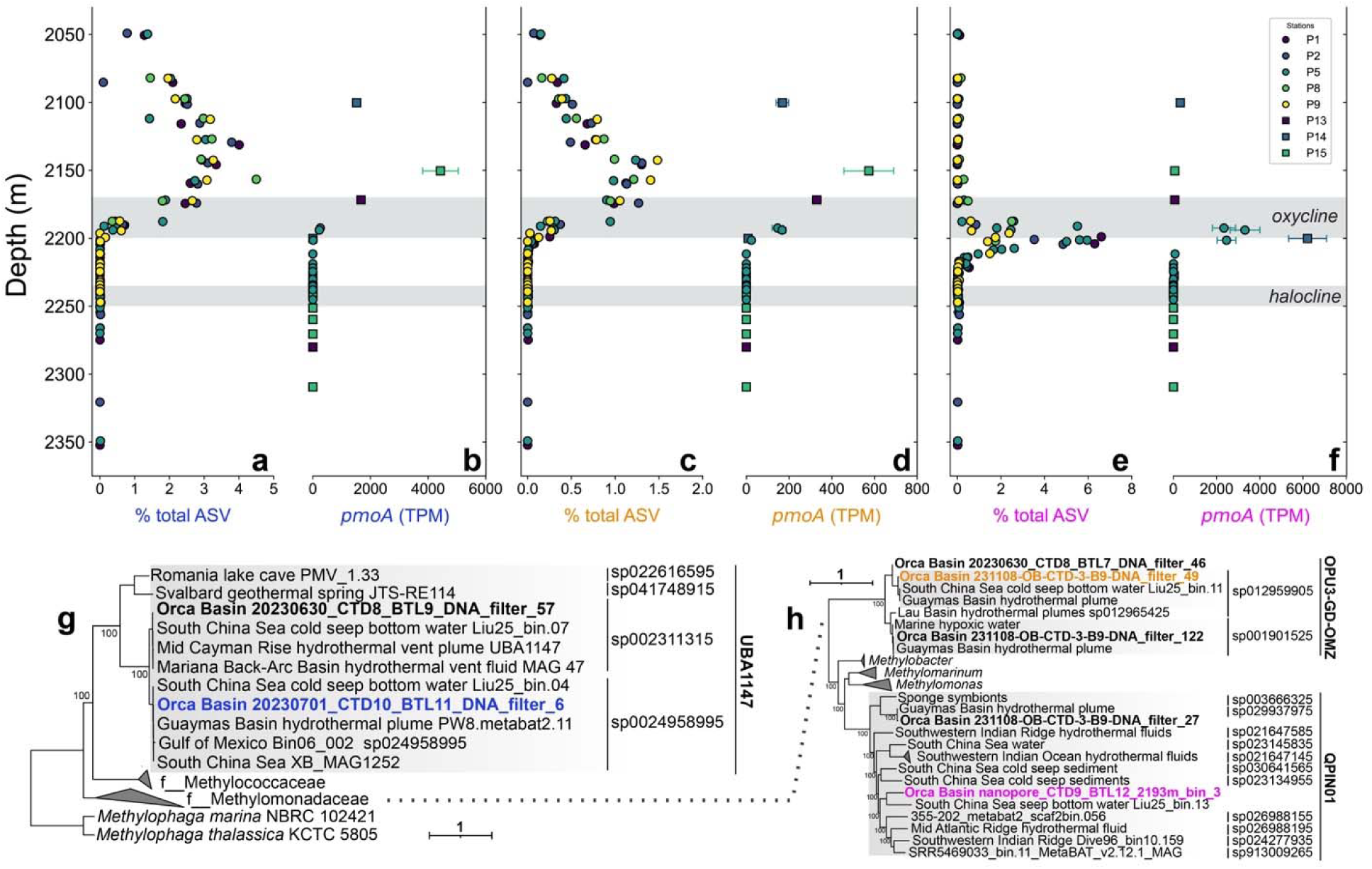
Relative abundance, *pmoA* transcription, and phylogeny of Methylococcales in Orca Basin. (**a**) Relative abundance (ASV %) of UBA1147; **(b)** *pmoA* gene transcription (TPM) of UBA1147; (**c**) relative abundance (ASV %) of OPU3-GD-OMZ; **(d)** *pmoA* gene transcription (TPM) of OPU3-GD-OMZ; (**e**) relative abundance (ASV %) of QPIN01; (f) *pmoA* gene transcription of OPU3-GD-OMZ; (**g**) phylogenetic tree of the UBA1147 family based on concatenated marker genes; (**h**) phylogenetic tree of OPU3-GD-OMZ and QPIN01 in the family Methylomonadaceae based on concatenated marker genes. In panels a-f, the upper gray box indicates the depth range of oxycline (2170-2200 meters) and the lower gray box indicates the depth range of the halocline (2235-2250 meters).

At the base of the oxycline (∼2200 m, <1 µM dissolved O_2_; **Table S1; Fig. 1c)**, genus QPIN01 (family Methylomonadaceae; MAG nanopore_CTD9_BTL12_2193m_3; 98.9% estimated completeness; **Table S2**), previously known as Marine Methylotrophic Group 2, sharply peaked in relative abundance (∼7% total ASV; **Fig. 5e**) and *pmoA* transcription (∼6000 TPM; **Fig. 5f**). QPIN01 has also been previously found to be abundant in other water columns with high methane flux, including above cold seeps [59, 60]. In Orca Basin, QPIN01 transcribed genes for low-oxygen adaption including dissimilatory nitrate reduction (*narG*), dissimilatory nitrite reduction (cytochrome cd-1 nitrite reductase (*nirS*)), and cytochrome bd-type oxygen reductase (*cydA*; **Table S1**) that are present in other aerobic methanotrophs that persist under oxygen limitation [58, 61, 62]. These genes were not identified in the other UBA1147 and OPU3-GD-OMZ MAGs, although these MAGs were incomplete **(Table S2**). Dissimilatory nitrate reduction by QPIN01 may contribute to the drop in NO_3_ ^-^ concentrations across the oxycline **(Fig. 1d)**.

## Conclusions

This study sheds new light on the microbial methane cycle of Orca Basin, particularly the methane biofilter overlying the brine, depicted in **Figure 6**. We corroborate previous findings that *Methanohalophilus* are the major methanogenic archaea in the NaCl-saturated brine (∼5 M Cl^-^) in Orca Basin [5, 6], as in other DHABs [10–15]. Our isotopic data indicates that methane from molecular diffusion out of the brine is anaerobically oxidized in the halocline at ∼2 M Cl^-^. Omics data suggest that ANME-2a, the most widespread and sequence-abundant ANME clade at cold seep sites [63], are the major driver of AOM in the Orca Basin. This finding was unexpected because the Orca Basin halocline salinity (2 M Cl^-1^) is double the previously known salinity maximum for ANME-2a of ∼1 M Cl^-1^ [31]. ANME-2a transcriptional activity was limited to a narrow depth range in the halocline; salt stress likely restricted AOM deeper in the water column, because the low energy yield of AOM does not generate sufficient ATP for energy-intensive adaptations to extreme salinity in the brine [17]. Long-term stability of the basin is likely important for the establishment of ANME populations, given their slow growth rates [64–66]. The capacity of the Orca Basin ANME-2a for osmolyte biosynthesis, apparently acquired from ANME-2d, may be key to its high activity in Orca Basin’s halocline. Identification of the bacterial partner of Orca Basin ANME-2a, possibly SURF-16/Seep-DBB, awaits further study. Whether Orca Basin ANME archaea ultimately depend on sulfate, manganese, iron, or another electron acceptor remains unclear. Sulfate is present at millimolar concentrations in the halocline and is sufficient to support conventional sulfate-AOM. Manganese oxide reduction is thermodynamically more favorable than sulfate and iron reduction [67, 68] and offers a plausible alternative or complement to sulfate-AOM in the particle-rich halocline. Additionally, the potential presence of Fe(OH)_3_ in the halocline may serve as another pathway complementary to sulfate-AOM and a recent report described ANME-2a-*Desulfobulbaceae* enrichments that may mediate Fe-AOM [47]. Sulfate-, Mn-, and Fe-AOM may therefore all contribute to methane oxidation in Orca Basin **(Fig. 6)**.

**Figure 6.**
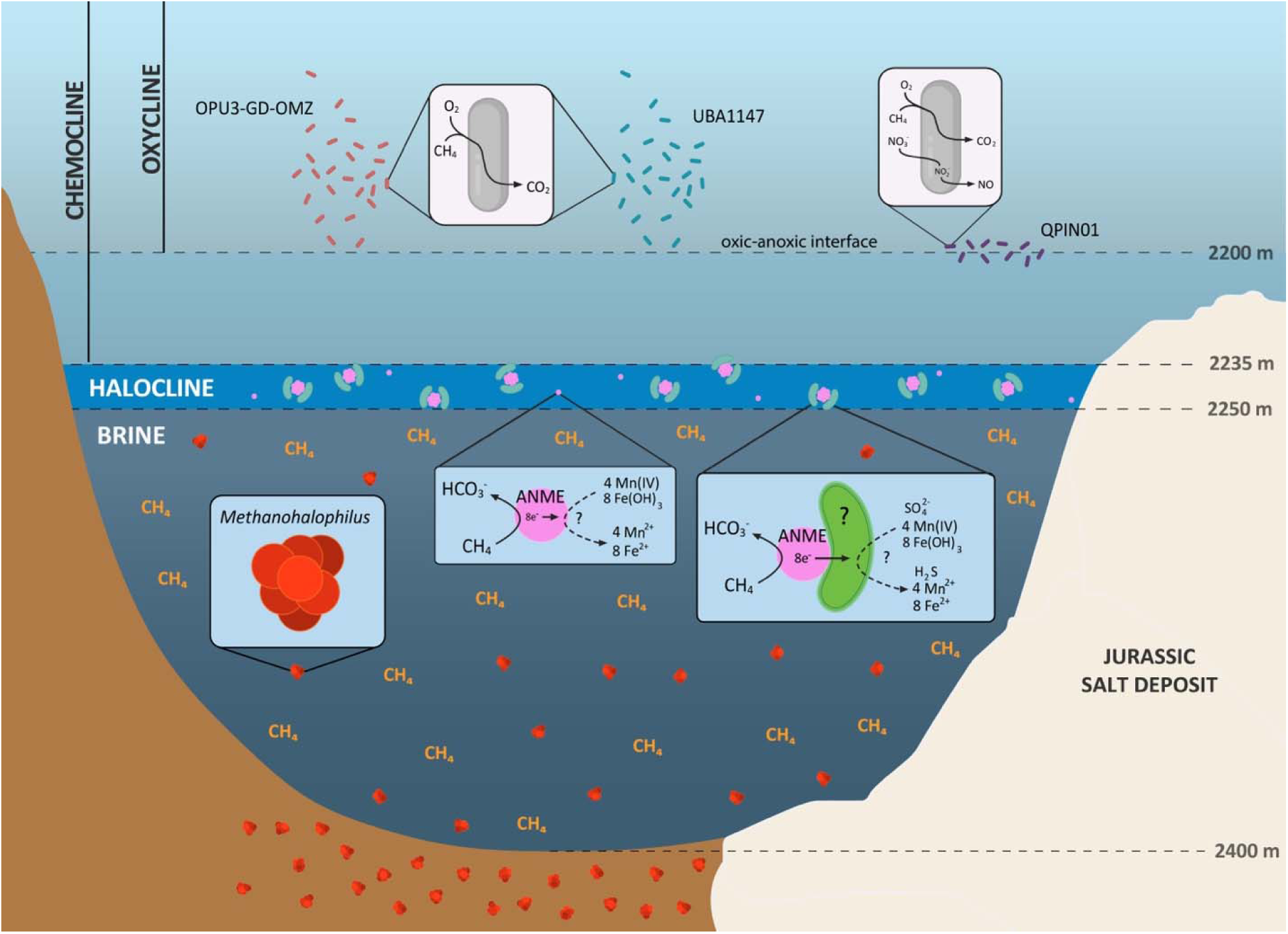
Hypothesized schematic of microbial methane cycling in Orca Basin. The chemocline between the overlying oxygenated seawater and methane-rich anoxic brine supports stratified microbial activity. In the oxic layer, aerobic methanotrophs (UBA1147, blue; OPU3-GD-OMZ, dark red; and QPIN01, purple) appear to oxidize methane. Within the anoxic brine, *Methanohalophilus* (red) may produce methane via methylotrophic pathways. Archaea (pink), likely to be anaerobic methanotrophic archaea (ANME), were observed forming consortia with unknown bacteria (green) in the particle layer, with an unknown electron acceptor. A portion of the ANME community may be free-living, performing Mn-AOM or Fe-AOM, independent of bacterial partners. The extent of the salt deposit is approximated from the cross-section in Pilcher and Blumstein [1].

Methane that escapes AOM encounters an oxygen-dependent biofilter mediated by aerobic Methylococcales bacteria in the Orca Basin oxycline **(Fig. 6)**. The genus QPIN01 is most transcriptionally active at the base of the oxycline (<1 µM dissolved O_2_) while genera UBA1147 and OPU3-GD-OMZ are active at 100-130 µM dissolved O_2_ in the oxycline **(Fig. 6)**. The metabolic flexibility of QPIN01, which appears to couple aerobic methane oxidation to partial denitrification, as previously reported for other methanotrophic gammaproteobacteria [58, 61, 62], may explain why this genus dominates at low oxygen concentrations while UBA1147 and OPU3-GD-OMZ are restricted to higher oxygen in overlying waters.

## Materials and Methods

### Sample collection and geochemical characterization

Samples spanning the transition from seawater (2050 m) to the brine core (2350 m) were collected by using Niskin bottles on a CTD rosette on 24 June-5 July 2023 from stations P1 (27.016°N, 91.274°W), P2 (27.027°N, 91.271°W), P5 (27.004°N, 91.251°W), P8 (26.908°N, 91.335°W), and P9 (26.919°N, 91.274°W) on the R/V *Point Sur* cruise PS23_26_Bowman, as described in Paris et al [23]. Additional samples were collected on 6-8 November 2023 from stations P13 (26.931°N, 91.335°W), P14 (26.927°N, 91.326°W), and P15 (26.919°N, 91.345°W) on the R/V *Point Sur* cruise PS24_13_Raven (OCRA-3). Detailed methods for geochemical characterization of stations P1, P2, P5, and P9 were previously reported in Paris et al [23], and data for P8 in this study was acquired in the same manner. Samples for sulfide concentration measurements were preserved in zinc acetate solution and subsequently quantified by the Cline assay [69].

### Methane concentrations and isotopic compositions

In June-July 2023, triple-rinsed syringes were used to collect seawater from each Niskin bottle immediately after rosette retrieval. Using gas-tight syringes, 30 mL of nitrogen gas was added to 30 mL of sample liquid for equilibration at room temperature by occasional shaking. After 4 hours, 12 mL of headspace gas was injected into an exetainer (Labco). The exetainers were stored in the dark for approximately two years. δ^13^C-CH_4_ was analyzed using a PreCon Automated Trace Gas Pre-Concentrator interfaced to a Delta V Plus isotope ratio mass spectrometer (Thermo Scientific) [70] at the UC Davis Stable Isotope Facility. Methane concentration measurements for samples collected on cruises 76-G-10, 77-G-2, 77-G-3, 77-G-13, and EN-32 were extracted from Wiesenburg et al [3] using WebPlotDigitizer. Depth offsets between the datasets are likely due to errors introduced by hand-graphing in the older dataset as well as differences in depth calibration from pressure data.

### DNA and RNA sample collection and extraction

Samples from a wide range of sampling locations and depths **(Table S1)** were collected for 16S rRNA, metagenomic, and metatranscriptomic sequencing. To obtain DNA samples for 16S rRNA and Illumina metagenomic sequencing, 1 L of seawater was collected from each depth, filtered using a vacuum manifold onto 0.2-μm 47-mm Supor-200 membrane filters (Pall), and stored at -80°C. To obtain RNA samples, triplicate 1-L water samples from each depth were immediately filtered onto 0.2-μm 47-mm Supor-200 membrane filters (Pall) using a Masterflex® peristaltic pump upon CTD recovery to minimize change of *in situ* gene expression. Each RNA sample was preserved in 5-mL cryovial with 1.5-mL of 0.2-μm filter-sterilized RNAlater at -80°C. DNA and RNA were extracted from each filter using the MagMAX Microbiome Ultra Nucleic Acid Isolation Kit and the KingFisher Flex high-throughput system (Thermo Fisher Scientific). RNA samples were treated with DNase to remove residual DNA using the TURBO DNA-free™ Kit (Invitrogen) according to the manufacturer’s instructions. DNA and RNA concentrations were quantified using the Qubit dsDNA and RNA High Sensitivity Assay (Invitrogen), respectively.

To obtain DNA samples for Nanopore sequencing, ∼4 L of seawater per depth was concentrated to 50-100 mL using a large-volume concentration kit (InnovaPrep CC01117) and further concentrated to 200-400 µL using a concentrating pipette (InnovaPrep CC08003-10, HC08018-P, HC08018-T). Eluent was stored at 4°C in DNA/RNA Shield (Zymo Research R1200). DNA was extracted using a ZymoBIOMICS DNA Microprep Kit (Zymo Research D4301) with the following protocol modifications: samples were agitated for 3 min with a TerraLyzer (Zymo Research S6022), centrifugation steps (16,000 x g) were repeated until supernatant was clear, samples were subjected to an additional 1-min dry spin to remove residual buffer after the final wash step, and samples were incubated for 3 min in 60°C DNase/RNase-free water before elution. II-μHRC Filter/HRC Prep Solution and III-F Filter steps were skipped. DNA concentration was quantified using the Qubit dsDNA High Sensitivity Assay (Invitrogen).

### 16S rRNA gene amplicon sequencing and analysis

16S rRNA gene amplicon sequencing was performed at the Argonne National Laboratory Environmental Sample Preparation and Sequencing Facility. Libraries were constructed with primers 515F and 806R [71] in a 151 x 151 architecture, and sequence processing followed the methods of [72]. Briefly, reads were quality controlled and denoised to amplicon sequence variants (ASVs) using dada2 prior to classification and metabolic inference with paprica [73], which uses Infernal [74], epa-ng [75], and gappa [76] to place ASVs onto a reference tree constructed from representatives in GenBank RefSeq [77].

### Illumina metagenomic sequencing and processing

Extracted DNA samples were submitted to the DOE Joint Genome Institute (JGI) for library preparation, sequencing, and downstream processing. Metagenome libraries were prepared by JGI using an Illumina low-input DNA protocol, sequenced on the Illumina NovaSeqX platform, and processed using JGI’s standard metagenome read filtering and quality control pipeline (SOP 1064.2). Raw reads were filtered and trimmed using the BBTools suite to remove adapter sequences, low-quality bases, homopolymers (homopolymers of G’s of ≥5 at the ends of the reads), optical duplicates, and reads failing minimum quality (<3 average quality score across the read) or length thresholds (≤51 bp or 33% of the full read length). Reads containing excessive ambiguous bases (≥4 ’N’ bases) were discarded. Potential contaminant-associated reads were identified by mapping against masked vertebrate reference genomes and removed, along with reads matching common microbial contaminants. The resulting high-quality reads were trimmed with BBtools bbduk.sh v39.01 using the following parameters deviating from default: <ref=adapters ktrim=r k=23 mink=11 hdist=1 tbo qtrim=r trimq=25 minlen=30>. Raw and trimmed read quality was checked with FastQC v0.11.8 (Andrews 2010). Trimmed reads were then assembled with megahit v1.2.9 on the <–meta-sensitive> preset, requiring a minimum contig length of 1500 bp. Bbtools stats.sh was used to assess assembly quality including N50. Trimmed reads were mapped back to assemblies to determine percentage of reads mapping using bowtie2 v2.5.1 [78] and SAMtools v1.15.1 [79]. Assembly was conducted separately for each sampling depth. SOURMASH v3.1.0 [80] was used to identify sample similarity at the nucleotide level. Samples of 15% or greater similarity were cross-mapped using bowtie2 v2.5.1 [78] and MAGs were generated using metaBAT2 v2 [81].

### Nanopore metagenomic sequencing and processing

Samples were purified and concentrated using the Genomic DNA Clean & Concentrator^TM^ Kit-10 (Zymo Research D4010). DNA (100-700 ng/sample) was barcoded and sequenced using a MinION R10.4.1 Flow Cell (FLO-MIN114) and the MinION Mk1C (Oxford Nanopore Technologies) following the “Ligation Sequencing gDNA with Native Barcoding Kit 24 v14” protocol (NBE_9169_v114_revP_15Sep2022; SQK-NBD114.24). Pod5 sequencing files were base called using dorado v0.8.0 (Oxford Nanopore Technologies) with the model dna_r10.4.1_e8.2_400bps_sup@v5.0.0 and separated into barcode-specific FASTQ and BAM files using the dorado demux command. Long-read fastq files for each depth were assembled and binned into MAGs with default parameters in Nanophase [82], which uses metaFlye [83], MetaBAT2 [81], MaxBin2 [84], MetaWRAP [85], SemiBin [86], minimap2 [87], Racon [88], and medaka (https://github.com/nanoporetech/medaka).

### Combined Illumina and nanopore MAG set analysis

Illumina and Nanopore MAGs were combined, and the resultant MAG set was assessed with CheckM v1.2.2 [89], dereplicated with dRep v3.4.5 [90], taxonomically classified with GTDB-tk v2.3.2 [91], and annotated with Prodigal v2.6.3 [92]. Code is available at https://github.com/rebeccasophiasalcedo/orca_master.

### Metatranscriptomic sequencing and analysis

All metatranscriptome libraries were processed by the DOE JGI following their standard metatranscriptome read filtering and quality control pipeline (SOP 1066.3). Libraries were prepared using an Illumina Ultra-Low Input RNA protocol and sequenced using the Illumina NovaSeqX platform. Raw reads were quality filtered and trimmed using the BBTools suite, including removal of adapter sequences, low-quality bases, homopolymers, optical duplicates, and reads below minimum quality or length thresholds. Reads containing ambiguous bases were discarded. Potential host-associated reads were identified by mapping against masked vertebrate reference genomes and removed, along with reads matching common microbial contaminants. Ribosomal RNA reads were removed using the BBTools RQCFilter workflow with JGI ribosomal k-mer references, and reads matching JGI metatranscriptome and KAPA spike-in reference sequences were identified and removed during the same filtering procedure. Removed rRNA and spike-in reads were written to dedicated, publicly available output files (JGI GOLD Study ID Gs0167160). The resulting high-quality, non-rRNA reads were mapped to predicted coding sequences for transcript quantification. A BWA index [93] was constructed on fasta files containing all coding sequences (CDS) identified by Prodigal [92] during MAG annotation. Metatranscriptome sequence reads were mapped to the index with BWA-MEM and SAMtools [79] used to identify mapped reads (-F 260). A custom script (https://github.com/bowmanlab/OAST/blob/main/count_mapped_reads.py) was used to count the number of reads mapped to each CDS for each sample, which were then converted to transcripts per million (TPM) [94].

### Characterization and phylogenetics of MAGs involved in Orca Basin methane cycle

MAGs containing marker genes for methanogenesis/AOM (*mcrA, mtrA*), aerobic methanotrophy (*pmoA*), and sulfate reduction (*dsrA*) were selected for further analysis. Multiheme cytochromes were screened using single_genome_multihemes.py (https://github.com/bondlab) and cell localization was predicted using PSORTb 3.0.3 [95]. An archaeal concatenated single-copy marker protein phylogeny of available Methanosarcinales MAGs (n=22, obtained from GenBank) was constructed using GTDB-Tk v2.7.2 [91] with the ar53 marker set along with the complete genomes for *Thermococcus* sp. 4557 and *Pyrococcus yayanosii* CH1 as outgroups. The phylogeny was created with FastTree 2.1.11 [96] in GTDB-Tk with the WAG model and CAT approximation with 20 rate categories. For the SURF-16 16S rRNA phylogeny, sequences were retrieved from NCBI BLASTN searches by querying the 16S rRNA sequence from the MAG nanopore_CTD9_BTL3_2236m_4 **(Table S2)**. Outgroup sequences were *Desulfonema limicola* 5ac10 and *Desulfococcus multivorans* DSM 2059. Sequences (n=32) were aligned using MAFFT online server with the L-INS-i method [97]. The maximum likelihood phylogeny was constructed in IQ-tree using model TPM3+I+R3 with ultrafast 1,000 bootstraps [98–100]. Bacterial concatenated single-copy marker protein phylogenies were also constructed using GTDB-Tk with bac120 marker set. As described in GTDB-Tk pipeline, maximum likelihood phylogenies were constructed with FastTree v2.1.10 [96] using the WAG model with 1,000 bootstraps [98–100]. All phylogenetic trees were visualized and annotated using iTOL [101]. Pairwise average nucleotide identity (ANI) of the Orca Basin *Methanohalophilus* MAG (231106-OB-CTD-1-B3-DNA_23; **Table S2**) to 8 *Methanohalophilus* reference genomes in Genome Taxonomy Database (GTDB-r232) was calculated with FastANI v1.34 [102] using default parameters.

### Catalyzed reporter deposition-fluorescence in situ hybridization (CARD-FISH)

Samples from 2234 m were subjected to a three-day anoxic incubation at 4°C with ^13^C-bicarbonate and ^15^N-ammonium as described in Paris et al [23]. Samples from a three-day incubation were used because cells were not fixed immediately after collection at this site/depth. Following incubations, cells were fixed with 2% formaldehyde for 12 to 18 hours at 4°C. Fixed cells were filtered under vacuum onto gold-coated polycarbonate filters, washed, dried, and stored at -20°C until analysis [103]. CARD-FISH hybridization reactions were performed at 46°C [104, 105] using ARCH915 (20% formamide) [106] and EUB338 (35% formamide) [107] oligonucleotide probes to target most archaea and bacteria, respectively. Cells were permeabilized with proteinase K, and tyramides conjugated to Alexa Fluor™ 546 (Thermo Fisher Scientific T20933) for archaea and Alexa Fluor™ 488 (B40953) for bacteria were used during amplification. Cells were stained with DAPI-based Vectashield (Vector Laboratories H-1200) and imaged with a Nikon ECLIPSE Ti2 microscope. Consortia were counted in 14 randomly selected fields of view (100 mm^2^ each) on a single filter **(Table S6)**. We also attempted CARD-FISH hybridization using the ANME-2a-647 probe [108] (50% formamide), which had no mismatches with the 16S rRNA sequence of the ANME-2a MAG (231106-OB-CTD-1-B3-DNA_filter_8; **Table S2**). The ANME-2a-647 probe did not yield interpretable signals under the conditions tested. The variable CARD-FISH performance of this probe has previously been noted [46].

## Data Availability

Metagenomic assembled genomes and 16S rRNA sequence data are available at NCBI BioProject PRJNA1380460. Metagenomic and metatranscriptomic data are available at DOE JGI GOLD study ID Gs0167160 and NCBI BioProject PRJNA1426066.

## Acknowledgements

We thank the captains, crew, and scientific teams of the R/V *Point Sur* (PS23_26_Bowman and PS24_13_Raven/Felker) for their support in sample collection and fieldwork. We thank Frederic Guilbault for the bathymetric map design. We thank Hang Yu for helpful discussions. This research was supported by the NASA Astrobiology Program (80NSSC18K1301; PI: B.E. Schmidt) and by the DOE Joint Genome Institute through the Community Science Program New Investigator Award (#510154) to E.J.S and A.M. Nanopore sequencing was conducted as part of work funded by NASA FINESST award 80NSSC22K1320 to J.M.M. and C.E.C. Sampling in 2024 (PS24_13_Raven/Felker) was supported by the Grantham Foundation for the Protection of the Environment (PI: M.R. Raven). We acknowledge the Stanford Computing Center and the Stanford Shared Geomicrobiology Facility (RRID:SCR_025000) for informatic and lab support. The Georgia Institute of Technology Jean “Chris” Purvis Endowed Professorship provided additional financial support.

