## Supplemental Figures for "Anaerobic methane oxidation by ANME-2a at 2 M chloride in the Orca Basin halocline"

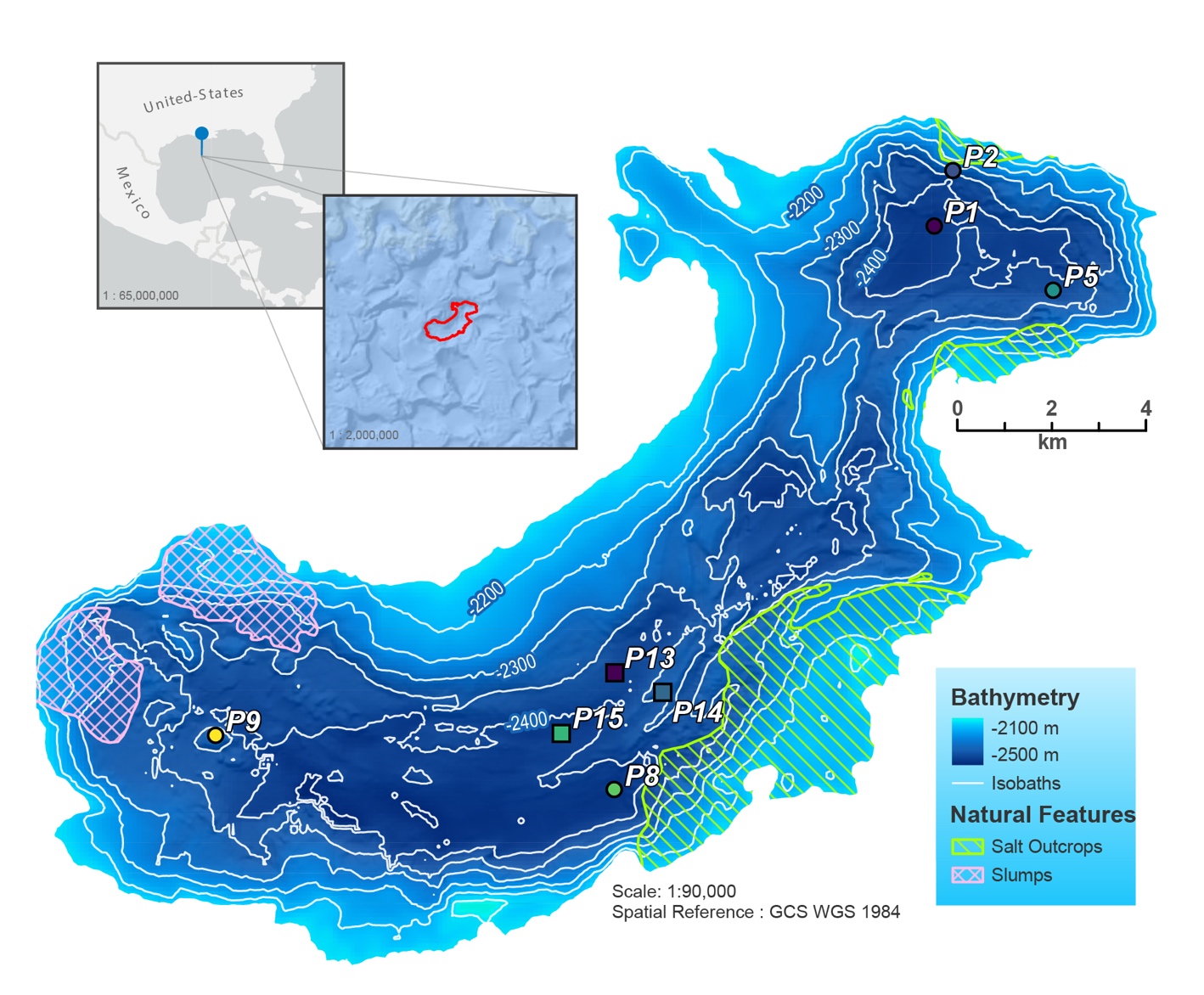

**Figure S1. Bathymetric map with sampling locations.** Bathymetry of the Gulf of Mexico along Texas-Louisiana continental slope (inset) with locations of sampling sites, modified from Paris et al [1].

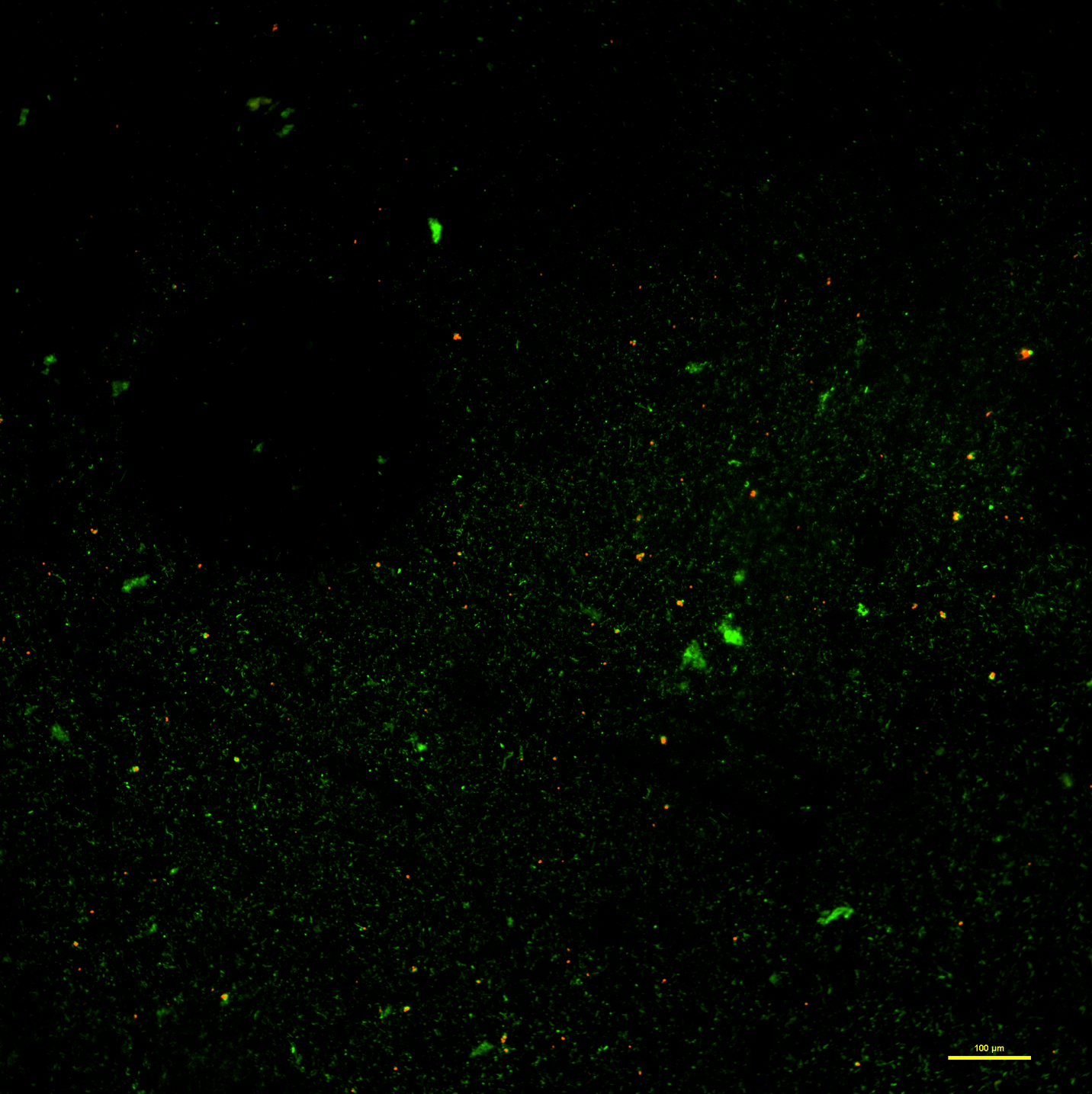

**Figure S2. CARD-FISH microscopy (10x magnification) of archaea-bacterial consortia in samples from 2234 meters depth** **in the Orca Basin halocline with probes ARCH915 (archaea; red) and EUB338 (bacteria; green).** Scale bar: 100 μm.

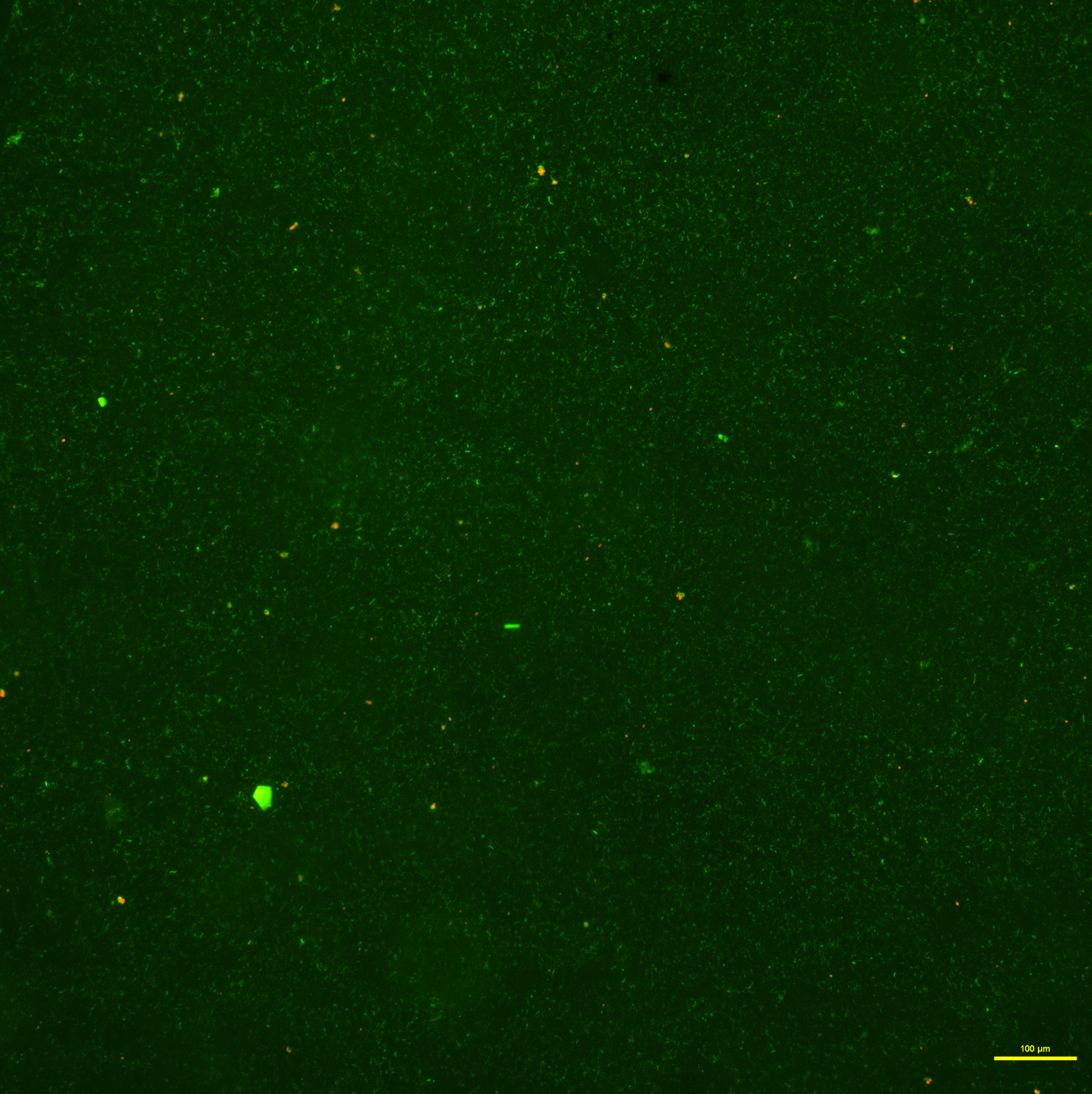

**Figure S3. CARD-FISH microscopy (10x magnification) of archaea-bacterial consortia in samples from 2234 meters depth** **in the Orca Basin halocline with probes ARCH915 (archaea; red) and EUB338 (bacteria; green).** Scale bar: 100 μm.

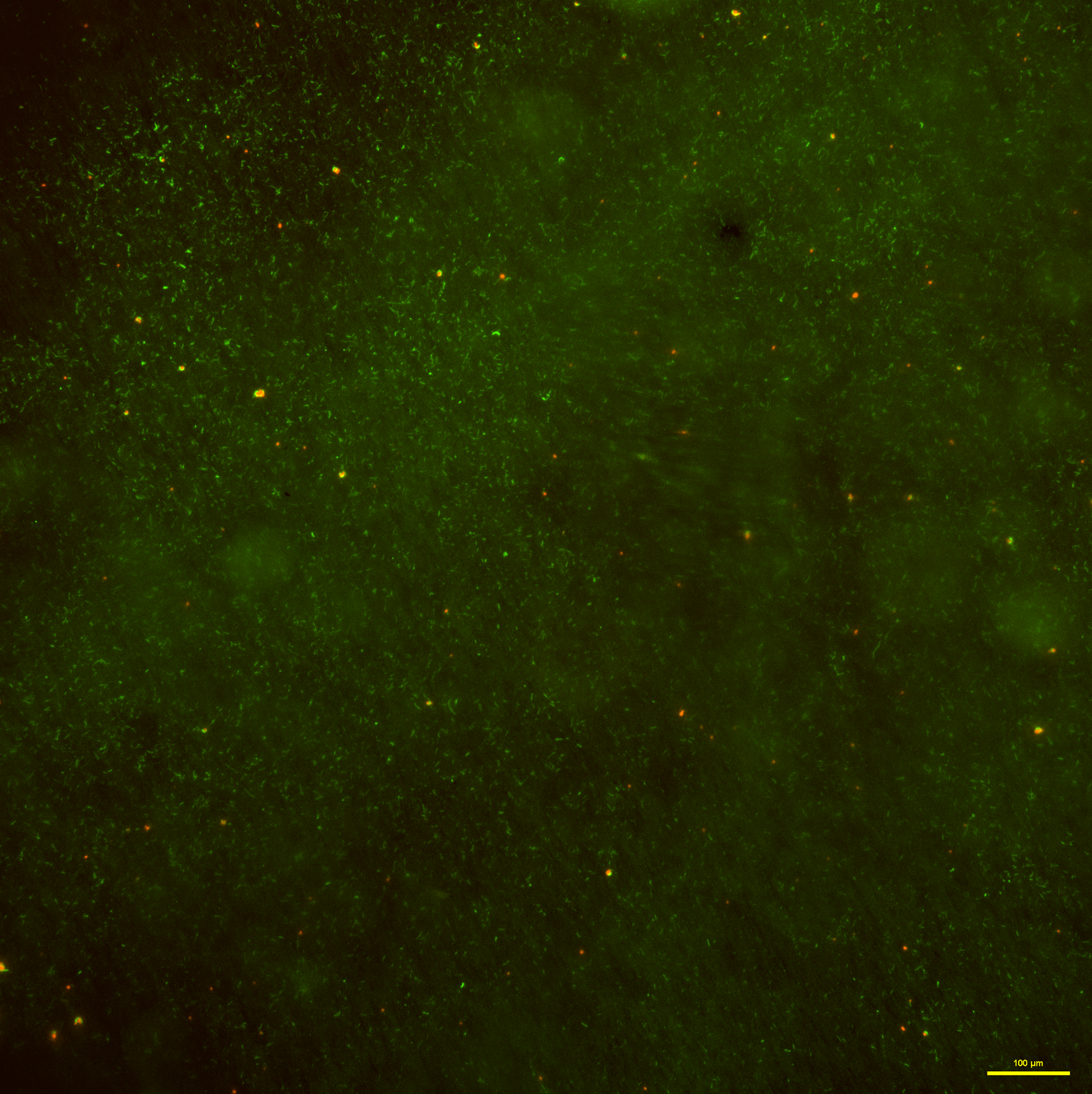

**Figure S4. CARD-FISH microscopy (10x magnification) of archaea-bacterial consortia in samples from 2234 meters depth** **in the Orca Basin halocline with probes ARCH915 (archaea; red) and EUB338 (bacteria; green).** Scale bar: 100 μm.

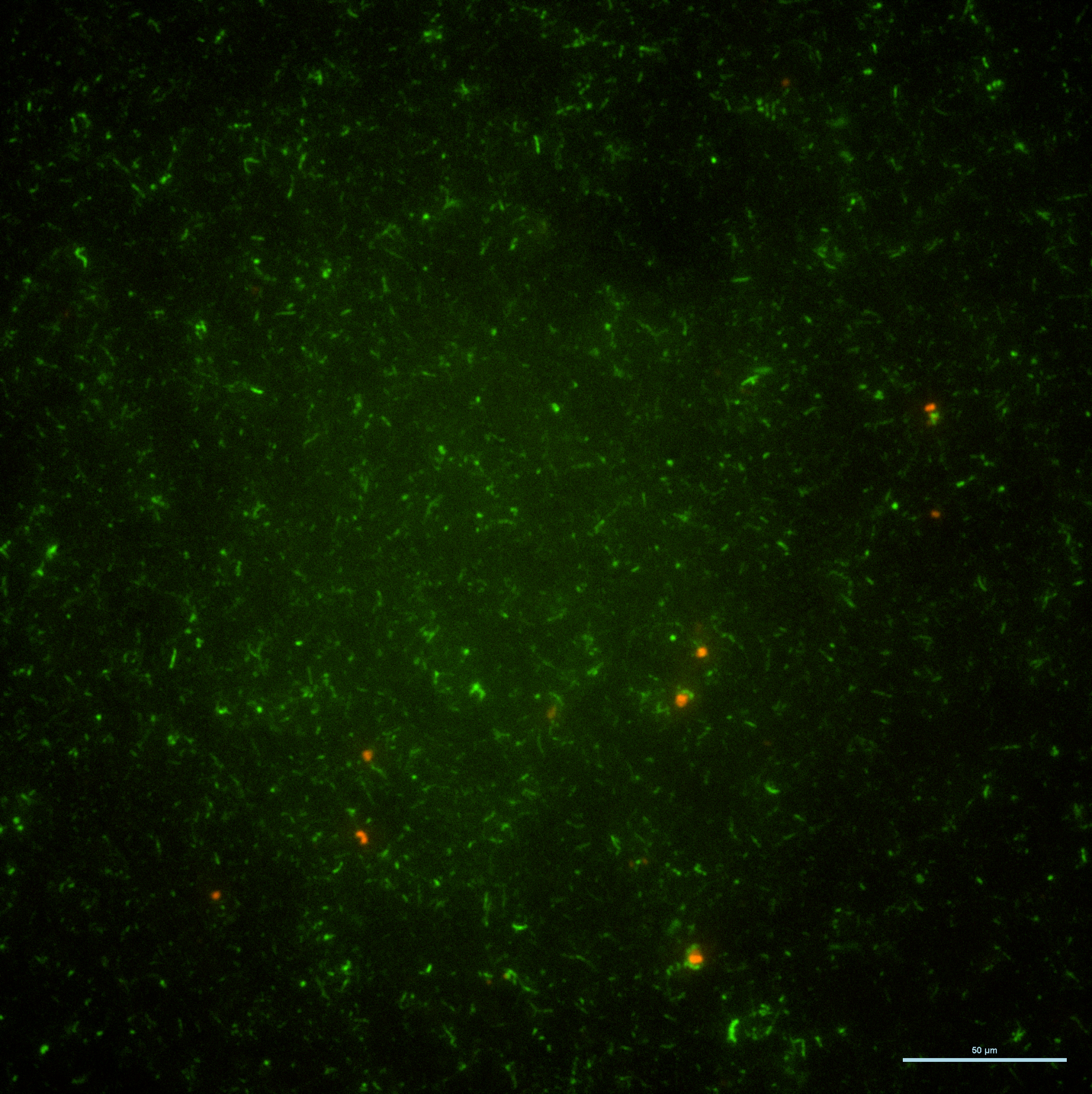

**Figure S5. CARD-FISH microscopy (40x magnification) of archaea-bacterial consortia in samples from 2234 meters depth** **in the Orca Basin halocline with probes ARCH915 (archaea; red) and EUB338 (bacteria; green).** Scale bar: 50 μm.

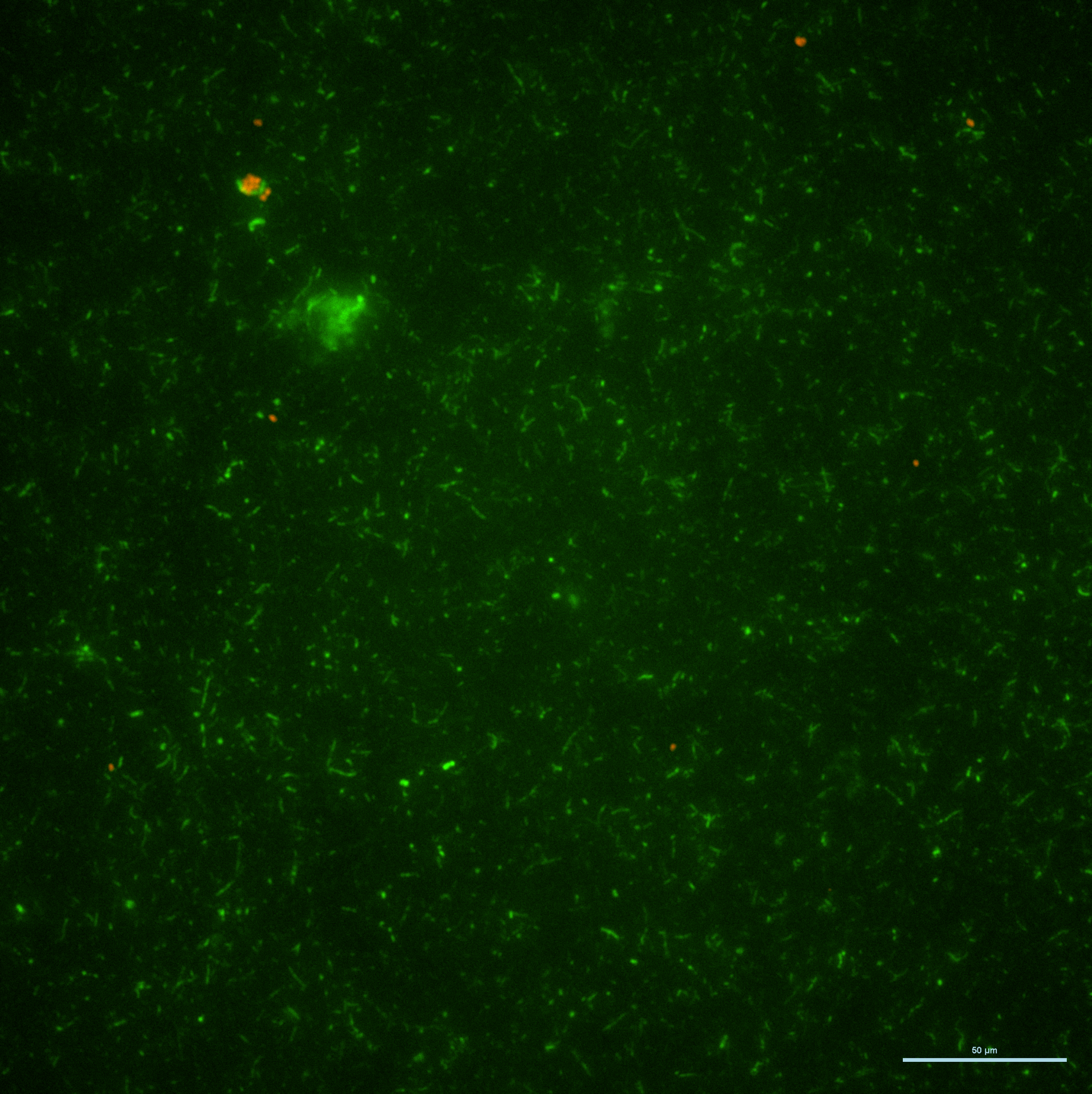

**Figure S6. CARD-FISH microscopy (40x magnification) of archaea-bacterial consortia in samples from 2234 meters depth** **in the Orca Basin halocline with probes ARCH915 (archaea; red) and EUB338 (bacteria; green).** Scale bar: 50 μm.

**Supplemental References**

1.	Paris ER, Salcedo RSR, Buessecker S *et al.* Free-living autotrophs drive a peak in carbon fixation in the Orca Basin chemocline while heterotrophs dominate total activity from seawater to brine. In revision after second review, ISME J
